# Construction of whole cell bacterial biosensors as an alternative environmental monitoring technology to detect naphthenic acids in oil sands process-affected water

**DOI:** 10.1101/2024.04.05.588297

**Authors:** Tyson Bookout, Steve Shideler, Evan Cooper, Kira Goff, John V Headley, Lisa M Gieg, Shawn Lewenza

## Abstract

After extraction of bitumen from oil sands deposits, the oil sand process-affected water (OSPW) is stored in tailings ponds. Naphthenic acids in tailings ponds have been identified as the primary contributor to toxicity to aquatic life. As an alternative to other analytical methods, here we identify bacterial genes induced after growth in naphthenic acids and use synthetic biology approaches to construct a panel of candidate biosensors for NA detection in water. The main promoters of interest were the *atuAR* promoters from a naphthenic acid degradation operon and upstream TetR regulator, the *marR* operon which includes a MarR regulator and downstream naphthenic acid resistance genes, and a hypothetical gene with a possible role in fatty acid biology. Promoters were printed and cloned as transcriptional *lux* reporter plasmids that were introduced into a tailings pond-derived *Pseudomonas* species. All candidate biosensor strains were tested for transcriptional responses to naphthenic acid mixtures and individual compounds. The three priority promoters respond in a dose-dependent manner, which allows semi-quantitative measurements, to simple, acyclic and complex NA mixtures, and each promoter has unique NA specificities. The limits of NA detection from the various NA mixtures ranged between 1.5 - 15 mg/L. The *atuA* and *marR* promoters also detected NA in small volumes of OSPW samples and were induced by extracts of the panel of OSPW samples. While biosensors have been constructed for other hydrocarbons, here we describe a biosensor approach that could be employed in environmental monitoring of naphthenic acids in oil sands mining wastewater.

## Introduction

The Athabasca oil sands in northern Alberta represent one of the world’s largest sources of recoverable bitumen (1). Bitumen is a heavily biodegraded crude oil recovered from surface mining the oil sands through the Clark process, or alkaline hot water extraction (2, 3). After separation of the oil from the extraction water, the resulting oil sands process-affected water (OSPW) is deposited and stored in tailings ponds where solids can settle, and the water can be recycled for repeat extractions (1). In addition to sand, silt, and clay, the OSPW contains many compounds such as dissolved ions, heavy metals, unrecovered oil, and numerous acid-extractable organic (AEO) compounds like naphthenic acids (1–3).

Naphthenic acids (NA) are naturally produced during the degradation of petroleum and are present in the oil sands ore used to produce bitumen. Naphthenic acid concentrations at the low end (5-30 mg/L) are in-line with concentrations observed in Athabasca wetlands (4), and mid-range concentrations (50 – 120 mg/L) are in-line with OSPW and industrially-affected experimental wetlands (3, 2, 5, 6). They are classically defined by the formula C_n_H_2n-z_O_2_, where n is the number of carbons and Z indicates the number of hydrogens lost due to ring formation, and comprise a complex mixture of monocyclic, polycyclic, and acyclic alkyl-substituted carboxylic acids (2). The NAFCs extracted from OSPW have been characterized at the molecular level by Headley et al. and others according to distribution of classical NA based on carbon number; double bond equivalence and number of rings; along with heteroatomic (e,g S, N) content (7, 4, 6). NA have been identified as the main contributor to OSPW toxicity, and have demonstrated toxicity in microbes, plants, fish, amphibians, birds and mammals (3, 5, 8). Some naphthenic acid compounds are difficult for bacteria to degrade, particularly those high molecular weight compounds containing multi-ringed structures and branched carboxylic acids (3).

Standard practice in the oil sands industry is to store OSPW in large tailings ponds. Despite OSPW recycling, this has resulted in the accumulation of over 1 billion m^3^ of OSPW (9). There are currently no established, cost-effective methods available to remediate the organic contaminants in the tailings ponds on the large scale required, prior to eventual discharge back to the natural environment. There are various analytical methods for monitoring and quantifying NA, including conventional screening techniques such as Gas Chromatography-Mass Spectrometry (GC-MS), Fourier Transform Infrared Spectroscopy (FTIR), along with molecular level high resolution mass spectrometry employing for example Fourier transform Ion cyclotron resonance or Orbitrap analysis with Negative Ion Electrospray Ionization-Mass Spectrometry (ESI-MS), with or without on-line High-Performance Liquid Chromatography (HPLC) (8, 10). These approaches are semi-quantitative with relatively low limits of detection, as low as 0.01 mg/L with ESI-MS (8). In general, the chemical analysis methods require multiple sample processing steps, including an organic solvent extraction of OSPW, which may bias the analysis, and can be time-consuming and costly (10). Hence, there is an opportunity for a rapid, cost-effective method for sensitive NA detection and quantification.

Whole cell biosensors are engineered bacterial strains capable of detecting and quantifying various compounds and analytes by producing a simple optical or electrochemical output proportional to the analyte of interest (11). This proven technology utilizes the refined metabolite sensing mechanisms of bacterial cells for detecting and adjusting to changes in the surrounding environment, providing a particularly useful tool for environmental monitoring. This technology is specific and capable of detecting low levels of small molecules, including aromatic compounds, alkanes, alkenes, heavy metals, and antibiotics (12–15). The sensitivity for biosensors are commonly in the parts per million range (mg/L), though there have been some reporting in the range of parts per billion (μg/L) (13, 16).

Bacterial biosensors are constructed by cloning a target analyte-induced promoter as a transcriptional fusion to a reporter gene (*lacZ*, *gfp*, *luxCDABE*), which allows for a simple, measurable output in response to exposure to the analyte in question. Using a *Pseudomonas species* OST1909 derived from a tailings pond (17), we performed RNA-seq to identify bacterial genes that are induced in response to various mixtures of naphthenic acids. We then printed a large collection of promoters and constructed a panel of transcriptional *luxCDABE* (bioluminescence) reporters to identify promoters that uniquely respond to acyclic naphthenic acids, a simple naphthenic acids mix, and the complex mixture of naphthenic acids that are extracted from OSPW, and untreated OSPW directly. NA-inducible promoters were configured in various biosensor designs, which were tested for their sensitivity and specificity of NA detection.

## Materials and Methods

### Library construction, SOLiD sequencing and RNA-Seq analysis

For RNA isolation, *Pseudomonas sp.* OST1909 (15) was grown (~25°C) in TB medium with 2% DMSO with or without 150 mg/L acyclic naphthenic acids (Sigma Aldrich 70340,15), or 150 mg/L of custom mix of 9X naphthenic acids (Table S1), or 150 mg/L of acid extracted organics; namely naphthenic acid fraction compounds (NAFC) extracted from oilsands process-affected water, which were prepared as previously described and provided by Environment and Climate Change Canada (19). TB is buffered media (~pH 7) was used to prevent any pH change upon addition of NA. Total bacterial RNA was isolated from duplicate, early-log cultures (OD_600_ = 0.2, 2×10^9^ cells) using the Qiagen RNeasy kit and stored in Qiagen RNA Protect. All subsequent steps were performed by the Centre for Health Genomics and Informatics at the University of Calgary.

Total RNA samples were treated with DNAse (Ambion) and assessed on an Agilent 2200 TapeStation. Then ribosomal depletion (Ribo-Zero, Illumina) and subsequent cDNA synthesis and conversion into sequencing libraries (SOLiD Total RNA-Seq Kit, Life Technologies) was performed as per the kit protocols. The indexed libraries were then pooled and added to SOLiD sequencing beads using emulsion PCR (EZ Bead E80 kit, Life Technologies). The resulting beads were then applied to two lanes of a six lane SOLiD 5500xl DNA sequencer for 75 bp sequencing. Each lane yield approximately 135 million reads (270M reads total). Transcriptome data has been assigned the GEO accession number GSE262045.

Reads from the RNA-Seq experiment were mapped to the *Pseudomonas sp.* OST1909 genome (17) using SHRiMP v2.2.3. (http://compbio.cs.toronto.edu/shrimp/; -v 50% -h 50%) (20). Samtools was then used to convert the .sam output into fasta format (http://www.htslib.org/) for transcript analysis with Kallisto v0.46.1-0 (https://pachterlab.github.io/kallisto/about, downgraded to legacy version that outputs bootstraps required by Sleuth (21) and the sleuth package in R (https://pachterlab.github.io/sleuth/about) (22). Kallisto quantified data by pseudo alignment, assigning reads to reference transcripts based on kmer counts (-k=31 -b 100 -s 1). Transcript abundances and bootstraps were modeled in Sleuth and subjected to statistical analysis via Wald test, modeled to remove transcripts occurring at low abundances across all conditions. The Mann-Whitney *U* test was performed to measure significant gene induction of all genes in an operon of interest.

### Construction of first, second and third-generation bioluminescent biosensors

A large panel of 54 promoters was targeted for DNA synthesis, combined with automated Gibson assembly to clone as transcriptional *luxCDABE* reporters in the plasmid pMS402 (23) (**Table S2**). The BioXP 3200™ was used for synthesis of predicted bacterial promoters, each with a minimum length of 165 bp that previously determined as a functional length for thousands of synthetic promoters (24). However, if no BPROM (25) predicted promoter was found within the first 165 bp upstream of the start codon, the sequence length was extended to include the BPROM predicted promoter. The synthesis was performed in a 96-well plate format which produced a recovery plate that contained 10 µl of transformation-ready ligated promoter *lux* constructs, in addition to 50 µl of each synthesized promoter sequence. The pMS402 reporter plasmid is first linearized by BamHI restriction digestion, and the promoter sequences are synthesized with an extra 30-40 bp on both 5’ and 3’ ends that are complimentary to the sequences flanking the BamHI site. The linearized vector and synthesized promoter were mixed with the Gibson assembly mix, which contains a 5’ exonuclease, DNA polymerase, and ligase enzymes. Except for linearizing the pMS402 vector, the cloning processes were conducted by CODEX DNA’s BioXP ™ 3200. For the first generation of synthesis and cloning, we maintained the pMS402 vector ribosome binding site (RBS) within 20 bp of the *luxC* start codon. Therefore, these constructs contained the promoter and vector ribosome binding sites (RBS) upstream of the *luxCDABE* operon. To optimize promoter cloning, we removed the vector RBS during the Gibson assembly in the second-generation constructs. The additional RBS between the promoter insertion site and the *lux* operon was hypothesized to interfere with expression of the other promoters. All plasmids were introduced into *Pseudomonas* OST1900 using electroporation.

A third-generation design was used in constructs with the *atuAR* and *marR* promoters, where we also included the corresponding regulator gene *atuR* or *marR*, respectively, upstream from the transcriptional reporter (**Fig 1**). The regulator gene was driven low strength (data not shown), constitutive OST1909 promoter upstream of a predicted BCCT family transporter (locus tag “IH404_RS26360”). To prevent gene expression read through, strong terminators (26) were added downstream of the two synthetic promoters (**Table S2**).

**Figure 1.**
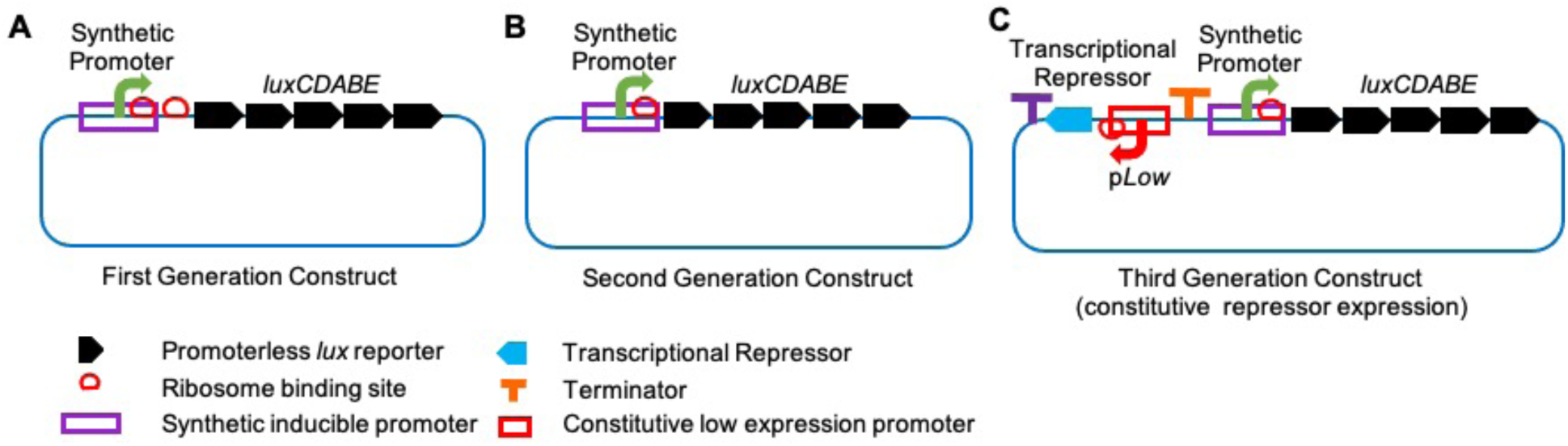
Genetic map of plasmid-based biosensor construct designs. **A.** The *luxCDABE* reporter genes (black) includes a RBS (red circle) upstream of the *luxC* start site. The first generation constructs include the target promoter cloned as a transcriptional *lux* fusion and includes a second RBS from the native promoter. **B.** The second generation constructs have the plasmid RBS removed during Gibson assembly. **C.** Third generation constructs also include the transcriptional repressor gene under the control of a constitutive, low level expression promoter. Transcriptional terminators were added to prevent any read through from upstream promoters.

### Mini-Tn*5*-*lux* mutagenesis of *Pseudomonas sp.* OST1909 and screening for NA-induced transcriptional fusions

As previously described (27), random mini-Tn5-lux mutagenesis was performed by mating the transposon from an *E. coli* donor into OST1909, and selecting insertion mutants on LB + 50 µg/mL tetracycline. Colonies were then stamped onto LB agar with and without 100 mg/ml of the custom mix of 9X naphthenic acids, and were visualized using a chemiluminescence imaging system, to identify colonies that had increased luminescence in the presence of NA. The genome of one mutant of interest was sequenced using minION (Oxford Nanopore) in order to map the *Tn* insertion site to the hypothetical gene with locus tag IH404_RS03680 (*hyp3680::lux*).

### High-throughput gene expression screening of biosensor constructs for specificity and sensitivity to naphthenic acids

Candidate biosensors were initially grown in tubes at room temperature (~25°C) overnight in 3 mL LB media + 50 µg/mL kanamycin, to maintain selective pressure on the promoter-*lux* plasmid and subcultured into M9 (Difco^TM^, 248510) or BM2 (0.5 mM Mg^2+^) (28) minimal, defined media with 20 mM succinate. The *lux* gene expression assays were conducted in black 96-well, clear bottom plates (Thermo Scientific). NA were diluted to their final concentrations in 99 µl growth media containing 2% DMSO to increase NA solubility, which was then inoculated with 1 µl of the overnight biosensor cultures. NA stock solutions were made according to **Table S1.** Other hydrocarbons were tested to ensure the biosensor specificity for NA and certain hydrophobic compounds such as alkanes and longer carbon chain carboxylic acids required the solvent of 45% ethanol and 55% polyethylene glycol 400 (PEGEt) (29), at a final concentration of 2%. A Breath-Easy® (Sigma-Aldrich) membrane was used to prevent evaporation during a 15-hour protocol in a PerkinElmer 1420 multilabel counter Victor^3^. The plate reader protocol (2 sec shake; gene expression in counts per second (CPS), growth (OD_600_)) included 45 time points, taken every 20 minutes. The gene expression CPS value was typically normalized by dividing the CPS by the OD_600_ for each read. To compare the bioluminescent response of the different biosensor strains to the various compounds tested, the Fold Gene Expression was also calculated by dividing the CPS/OD_600_ values of the NA treated sample by those of the untreated sample. A fold change of 1 indicates no change, and a fold change of 2 indicates a doubling of *lux* expression in comparison to the untreated control.

### Oil sands process water (OSPW) sample testing

Water samples from the tailings ponds in the mineable oil sands region, as well as naphthenic acid fraction components (NAFC) extracted from the OSPW, were provided by Environment and Climate Change Canada. Details are given elsewhere of the collection of OSPW; preparation of the NAFC extracts, and molecular level analysis of NAFC by negative-ion electrospray ionization high resolution Orbitrap mass spectrometry (4, 19). The concentrations of NAFC in both the water samples and extracts were determined and recorded in **Table S1**. Depending on the reported concentrations, the NAFC extracts were diluted by a factor of 10 or 100 and tested at a final concentration between 20 and 60 mg/L. To monitor NA levels in the water samples with minimal treatment or dilution, 90 μl of each sample were added to each well of the assay plate. This was supplemented with 10 μl of 10x BM2 to allow for biosensor growth, inoculated with 1 μl biosensor culture, and measured as described above.

## Results and Discussion

### Transcriptome analysis of genes induced during growth in the presence of naphthenic acids

In the presence of petroleum hydrocarbons, transcriptome studies revealed that bacterial genes required to transport, metabolize or defend against these environmental compounds are frequently induced (30–32).To identify genes regulated by exposure to naphthenic acids, we performed RNA-seq experiments with *Pseudomonas sp*. OST1909 (17) grown to mid-log phase in growth medium containing naphthenic acids. To generate a diverse array of NA-induced genes, transcriptome experiments were performed after adding 150 mg/L of three distinct naphthenic acid mixtures: acyclic naphthenic acids (15) (Sigma-Aldrich, 70340), a custom mix of 9 individual naphthenic acids (**Table S1**), or NAFC extracted from oilsands process-affected water (4, 19).

The upregulated genes included many genes annotated in metabolisms, fatty acid degradation, nutrient transport in and out of the cell, and here we focussed on upregulated metabolite sensing, transcriptional regulators (12) and their adjacent operons (**Table S3**). RNA-seq showed significant upregulation of genes under experimental conditions: OSPW (p<0.05 = 56; q<0.05 = 0; n=2), acyclic NA (p<0.05 = 140; q<0.05 = 18, n=2), and 9xNA (p<0.05 = 110; q<0.05 = 14, n=2). The genes induced by the custom 9xNA mix involved in fatty acid degradation (β-oxidation) include the *fadAB* genes (IH404_RS09140/5), *fadD1* (IH404_RS22010), *fadL* (IH404_RS20310) (33). The genes induced by the acyclic NA mixture that encodes another possible NA degradation operon is the IH404_RS15470-IH404_RS15515 cluster (**Table S3**). We prioritized 54 NA-induced promoters for construction as plasmid-based transcriptional fusions, as candidates for bacterial biosensors to be used to detect naphthenic acids (**Table S2**) based on their upregulation in the transcriptome, and a predicted role in naphthenic acid degradation, efflux, transport and small molecule sensing transcriptional repressors.

### Rapid construction of plasmid-encoded transcriptional fusions of NA-inducible promoters controlling the *luxCDABE* transcriptional reporter genes

Among the 54 promoters identified by RNA-seq, 26 promoters were induced in response to a commercially available mixture of acyclic carboxylic acids, 14 promoters in response to the simple mixture of nine NA compounds, 14 promoters in response to the naphthenic acids that were acid extracted from OSPW (**Table S2, S3**). For large scale printing of bacterial promoters, we used short upstream regions (165-350 bp) that contained predicted sigma-70 promoter sequences (25) to drive the *luxCDABE* reporter. The first-generation constructs included the ribosome binding site (RBS) within the promoter, in addition to a second RBS present in pMS402 upstream of the *luxC* start codon. In the second-generation constructs, we removed the vector-encoded RBS and included only the RBS present in the predicted promoter region from *Pseudomonas sp*. 1909 (**Fig 1**).

### The *atuA* and *atuR* promoters from the *atu* operon are induced after detection of acyclic naphthenic acids

After a preliminary screen of all candidate biosensor strains, we selected the mostly highly induced promoter from each of the three NA mixtures for further study. In response to the acyclic NA mixture, we identified an operon that encodes homologs of the *Pseudomonas aeruginosa* acyclic terpene utilization operon (*atu* operon) (34–36) (**Fig 2, Table S3**). While individual genes within the cluster were induced, we also confirmed induction of the whole *atu* operon using the *U* test (p<0.05, **Table 1**). In *P. aeruginosa*, terpenes and the carboxylic acids citronellate and geranylate are degraded by the *atu* genes, β-oxidation and the leucine-isovalerate utilization (*liu*) pathways (34–36). Notably, the *P. aeruginosa atu* operon is induced in the presence of citronellate and the acyclic monoterpenes citronellol and geraniol (34–36). The probable *Pseudomonas sp*. OST1909 NA biodegradation cluster *atuA* through *atuF* (IH404_RS19870-IH404_RS1990) is divergently expressed from the tetR-type transcription regulator AtuR (IH404_RS19865) (37) (**Fig 2A**). Divergent promoters were detected in this intergenic region (**Fig 2C**) and therefore both promoters were cloned as transcriptional *lux* reporters.

**Figure 2.**
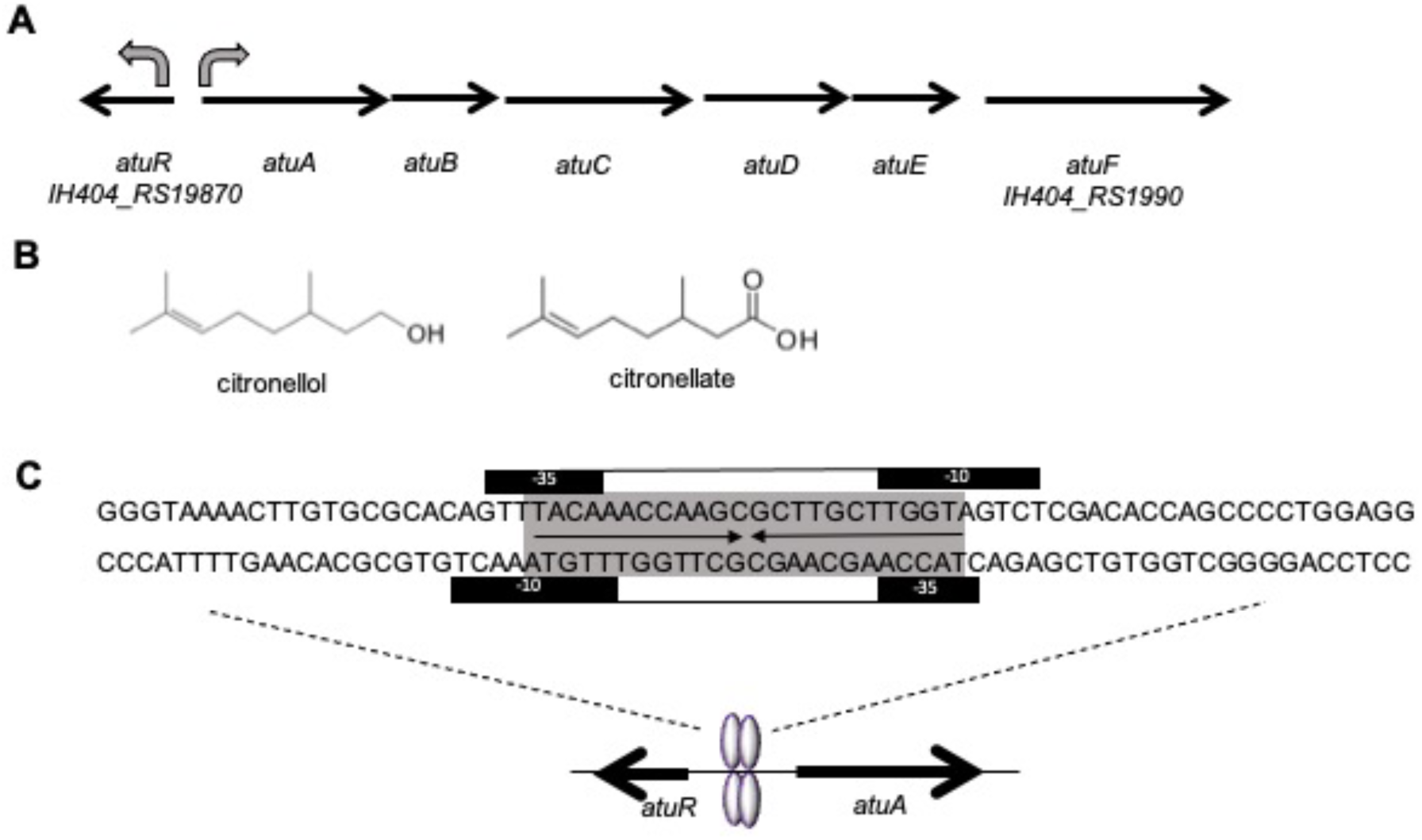
Gene and promoter map of the *Pseudomonas sp*. OST1909 *atu* operon. **A.** Arrow diagrams demonstrate the order and orientation of the *atu* genes within the operon. Sigma70-like promoters were found in the forward and reverse strand of the intergenic region between *atuR* and *atuA*, shown as curved arrows. **B.** Chemical structure of citronellol (acyclic terpene) and citronellate, which is a branched, acyclic naphthenic acid (or terpene). **C.** Predicted, overlapping sigma-70 promoters with −10 and −35 regions. AtuR dimers likely bind to inverted repeats (shown in grey) and represses expression of both divergent promoters.

**Table 1.**
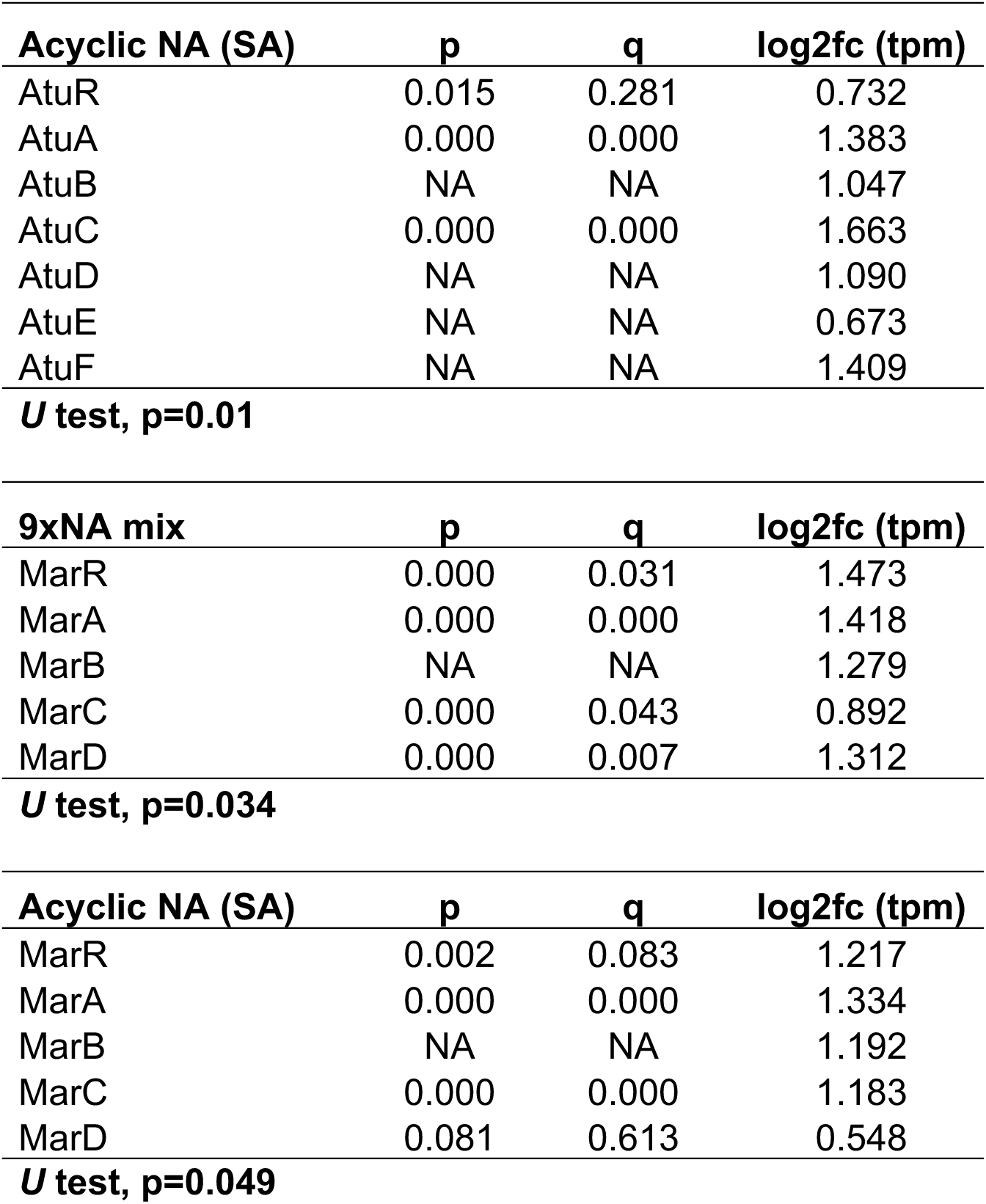
*U* test analysis of upregulated operons from RNA-seq data.

Based on the model of the *P. aeruginosa* AtuR repressor (37), a homolog of the TetR repressor, we predicted that the *Pseudomonas sp*. OST1909 AtuR homolog binds to and therefore senses acyclic naphthenic acid compounds, which relieves this repressor from its inverted repeat binding site, resulting in induction of the *atuA-F* cluster in the presence of naphthenic acids. In *Pseudomonas sp*. OST1909, the *atu* cluster is induced by acyclic naphthenic acids, which are likely degraded by these genes, which are similar in structure to acyclic terpenes (**Fig 2B**).

As shown in **Figure 3**, both first and second generation p*atuA*-based biosensors display a strong bioluminescent response to the acyclic NA mixture in defined M9 media. For this comparison experiment, we used relatively high naphthenic acid concentrations of 400 mg/L. In contrast to the complex NA mixtures extracted from OSPW, this commercially available naphthenic acid extract is from a poorly defined petroleum source, and mass spectrometry analyses of this sample demonstrated that most compounds are acyclic naphthenic acids and range in length from 5-20 carbons (38, 18). Expression of most biosensors was compared in BM2 and M9, both of which are similar defined growth media, and overall, the base level expression and the naphthenic acid induction responses were generally higher in M9 (data not shown). While the *atuA* expression levels are generally lower in the first-generation biosensors, the peak fold gene expression in the first-generation *patuA^1^-lux* biosensor is higher (~8 fold) than that of the second-generation sensor (~3.5 fold) (**Fig 3A**). In general, we interpret the biosensors with highest induction response (peak fold change) as the most sensitive strains in response to the analytes tested. The lower gene expression of the first-generation biosensor may be attributed to the extra 31 bp of DNA and the second RBS found within, which may interfere with the translation of the *lux* operon.

**Figure 3.**
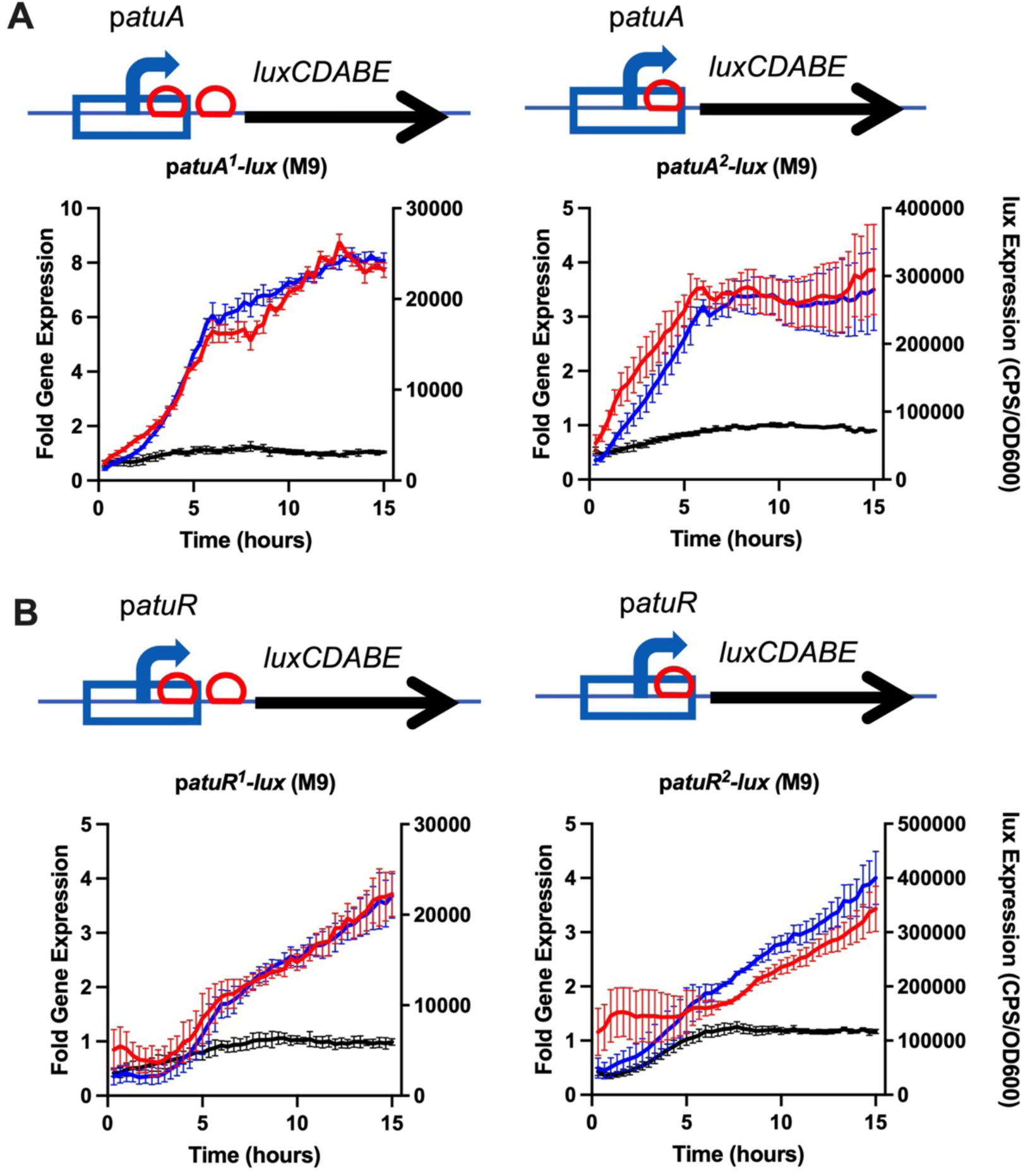
Gene expression response of first and second generation *atuA* and atuR-based biosensors to the acyclic NA mix. **A)** First and second generation *patuA-lux* biosensors and **B)** first and second generation *patuR-lux* biosensors were cultured in M9 medium with 400 mg/L acyclic mix and *lux* expression (CPS/OD_600_) over time was compared to untreated controls, shown in blue and black, respectively. The fold induction (shown in red) is calculated by dividing the expression in the presence of NA by the control media. All values shown are the average and standard deviation of triplicates, and each experiment was performed 3 times.

### The *atuR* promoter is also induced in response acyclic naphthenic acids

The divergent *atuA* and *atuR* promoters are predicted to be regulated by AtuR, a transcriptional repressor from the TetR family of regulators (39, 40) (**Fig 2**). Binding to the inverted repeat operator by AtuR would repress both promoters, and both promoters are predicted to be induced by naphthenic acid binding to AtuR. The *atuR* promoter was also cloned as first and second-generation biosensors. While the second generation *patuR^2^-lux* had roughly a 10X fold higher levels of overall expression, there was very little difference in the maximum fold gene expression (4-fold) between the two biosensor designs, which might be due to the slower induction response of this promoter compared to p*atuA* (**Fig 3B**). The p*atuR*-based biosensor construct reaches a maximum fold induction after 15 hours, compared to the maximum response time of around 5-7 hours for the p*atuA*-based construct (**Fig 3**). In conclusion, the *atuA* promoter responds faster and to higher fold changes to acyclic naphthenic acids, compared to the *atuR* promoter, and is therefore a better choice as for biosensors that detect acyclic naphthenic acids. It should be noted that gene expression responses to naphthenic acids begin within minutes, although the maximal responses are within hours, indicating a very fast sensing bacterial response (**Fig 3**).

### The *atuA* promoter responds specifically to acyclic naphthenic acids and not to related hydrocarbons

To test the range of specificity, the atuA-L biosensor construct (see methods for details) was screened with decreasing concentrations of four different NA mixtures, as well as 28 different individual hydrocarbon compounds at a concentration of 50 mg/L. **Figure 4** indicates the specificity of the atuA-L biosensor is primarily to acyclic naphthenic acids. When exposed to diverse NA mixtures, the atuA-L construct displays significant 2-fold *lux* expression patterns for both the commercially available acyclic NA mixtures (18) (Sigma-Aldrich, Acros) at 50 mg/L (**Fig 4A**). The individual compounds inducing the highest *lux* response from atuA-L are the acyclic NA compounds with low and medium length carbon chains such as hexanoic acid (2-fold), citronellate (3-fold), pentadecanoic acid (2.5-fold), and stearic acid (~2.8-fold) (**Fig 4B**). The *atuA* promoter does not respond to any other related compounds, including ringed naphthenic acids, alkanes or BTEX (benzene, toluene, ethylbenzene, xylene). Although the *atu* operon is annotated in *P. aeruginosa* for the utilization of acyclic terpenes (citronellol) and citronellate (16), the atuA-L biosensor responds to citronellate but does not exhibit a bioluminescent response to terpenes like citronellol or geraniol (**Fig 4B**). This suggests that the oil sands bacterial isolate *Pseudomonas sp.* OST1909 uniquely induces the *atu* operon in response to acyclic naphthenic acids, rather than acyclic terpenes.

**Figure 4.**
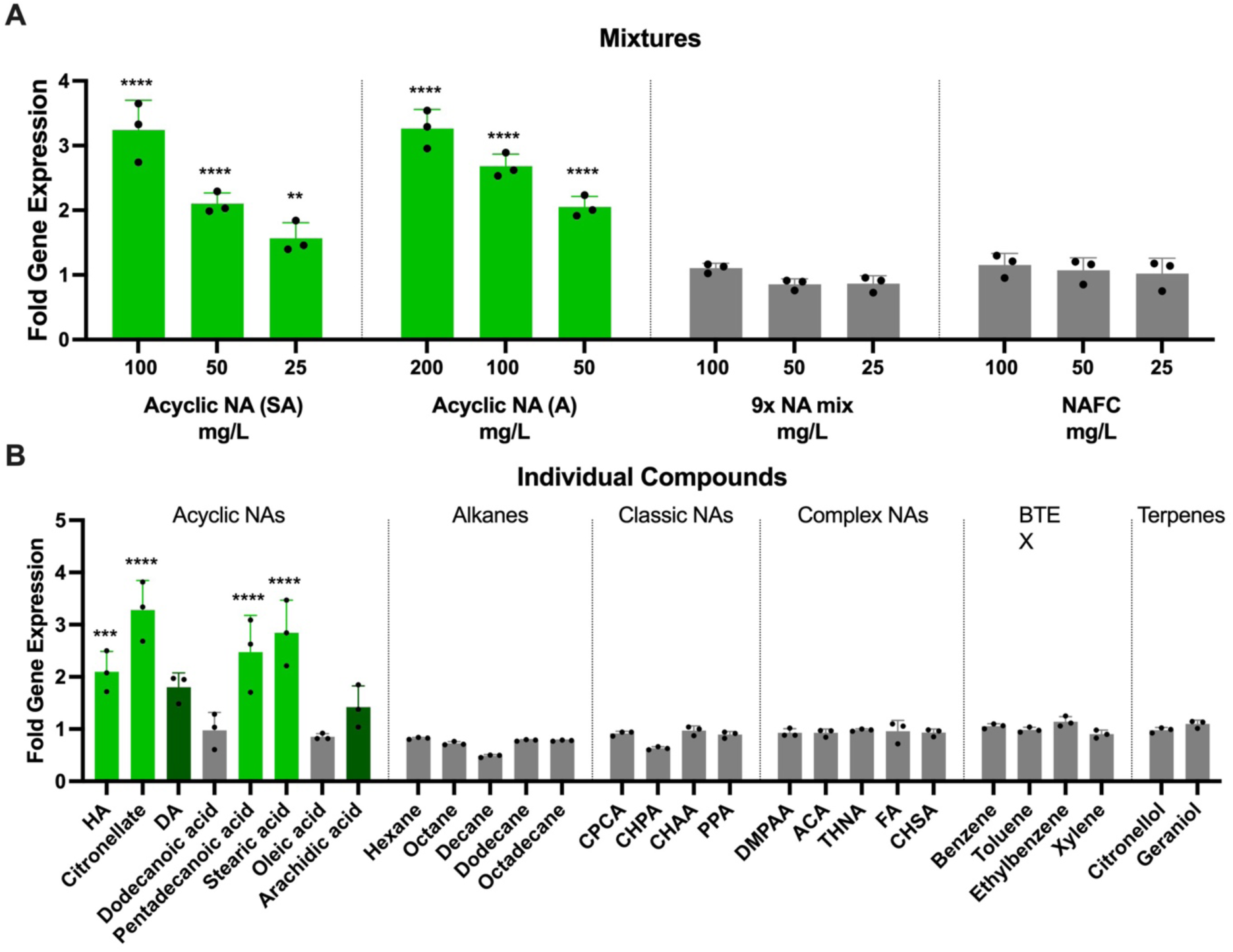
Specificity of the atuA-L biosensor to naphthenic acid mixtures and individual compounds. **A.** The atuA-L biosensor was exposed to four different naphthenic acid mixes with decreasing concentrations to monitor Fold gene expression values. All values represent the average and standard deviation of triplicate values taken at 6.6 hr. Sigma-Aldrich (SA) and Acros (A) indicate two commercial sources of acyclic NA mixtures. A two-way ANOVA found that there was a statistically significant difference between the mixtures (F(3,32) = 134.6, p < 0.0001) and concentrations (F(3,32) = 81.12, p < 0.0001). **B.** Fold gene expression values of 28 different hydrocarbons at a concentration of 50 mg/L. All values represent the average and standard deviation of triplicate values taken at 3.3 hours. A one-way ANOVA revealed a statistically significant difference between treatment groups (F(28,58) = 20.96, p < 0.0001). A post hoc Tukey HSD test was then conducted with each compound and mix to test significant relationships between each group (α = 0.05, n = 3). Treatment groups with significant results compared to the control are indicated with asterisks and bright green (*** indicate p<0.001, **** indicate p<0.0001). Nonsignificant groups displaying fold gene expression values above 1.5 are shown in dark green.

### The promoter regulating *marR* and antimicrobial resistance genes responds to fusaric acid, an antibiotic with a naphthenic acid structure

Among the genes identified from the transcriptome study, an operon containing a MarR family regulator (41) and genes predicted to contribute to fusaric acid transport, efflux and resistance (**Fig 5**), was found to be upregulated in the transcriptome study by the acyclic NA mixture (p<0.05, **Table 1, S3**). Fusaric acid is an antibiotic with a naphthenic acid structure that has an aromatic ring containing a nitrogen heteroatom and two hydrocarbon branches (**Fig 5B**).

**Figure 5.**
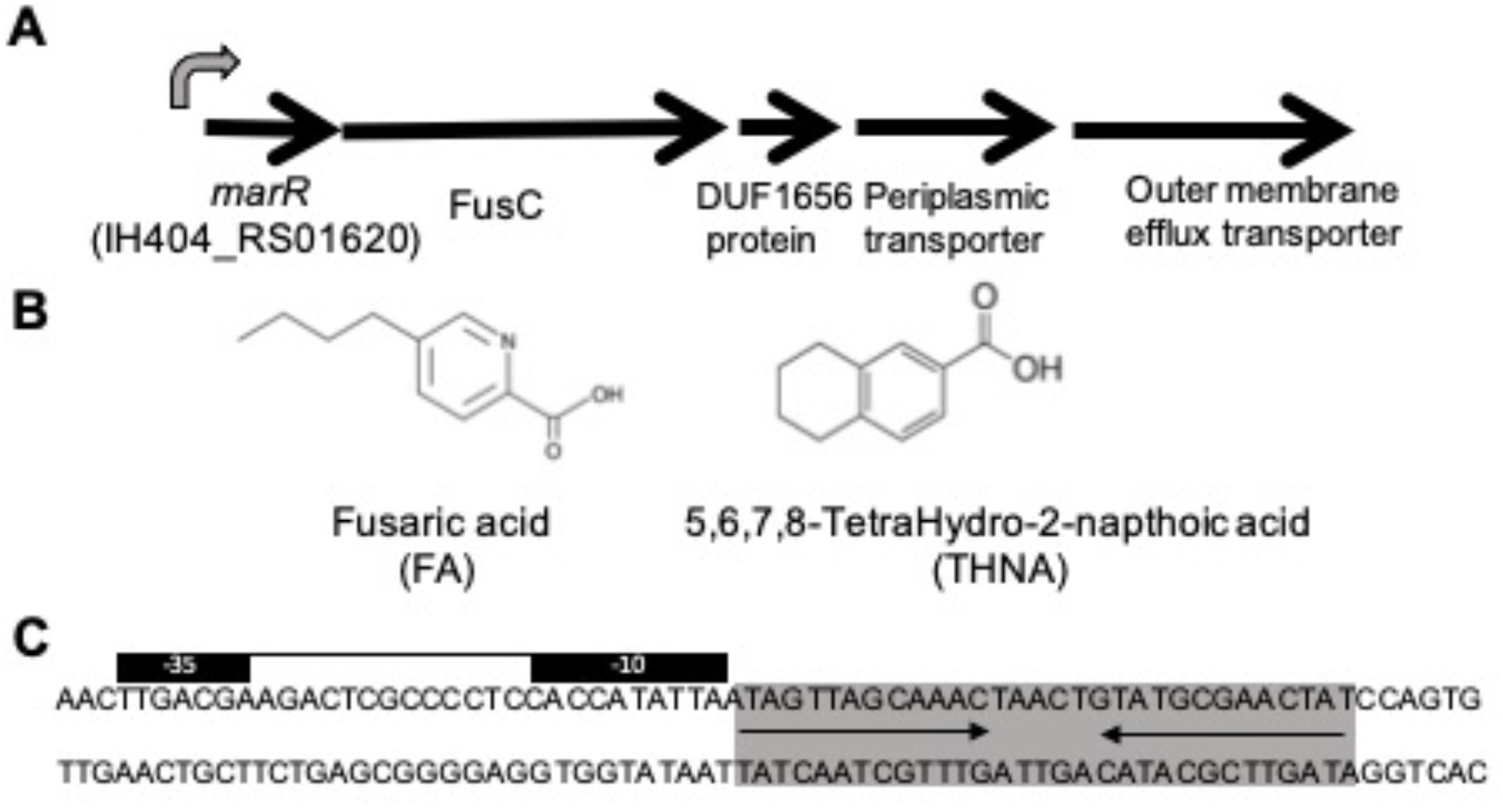
Gene map and promoter of the *marR* and naphthenic acid resistance operon. **A.** Arrow diagrams demonstrate the order and orientation of the genes encoding a MarR family transcriptional regulator, FusC fusaric acid resistance transporter protein, and an RND efflux pump. **B.** Fusaric acid has a naphthenic acid-like chemical structure, and THNA is another complex naphthenic acid that induces *marR* expression **C.** The sigma70-like promoter is located upstream of *marR*, indicated by the −10 and −35 regions, which would be repressed by MarR binding to the predicted inverted repeat binding site downstream of the promoter.

The *marR* gene found in *Pseudomonas sp.* OST1909 encodes a regulator protein that is part of a large and widely distributed family of <u>m</u>ultiple <u>a</u>ntibiotic <u>r</u>esistance regulators (MarR) (41). This family of transcription factors is conceptually similar to the TetR family, controlling many other cellular processes such as antibiotic resistance, stress response (42), virulence (43), and degradation or export of harmful compounds (44). Using the promoter region found upstream of the *marR* operon, first and second generation p*marR-lux* biosensors were constructed and tested for their responses to multiple naphthenic acid mixtures. We demonstrated that both p*marR*-based biosensor constructs were strongly induced by fusaric acid, an antimicrobial with an NA-like compound with a nitrogen-containing ring structure (**Fig 5B**). Similar to the p*atuA*-based biosensors, the first generation *marR* sensor has lower overall expression than the second generation biosensor, but higher induction responses to fusaric acid, therefore greater sensitivity (**Fig 6**). Gene expression reached a maximum within 3 hours and rapidly decreased thereafter, which may represent a pattern of negative autoregulation.

**Figure 6.**
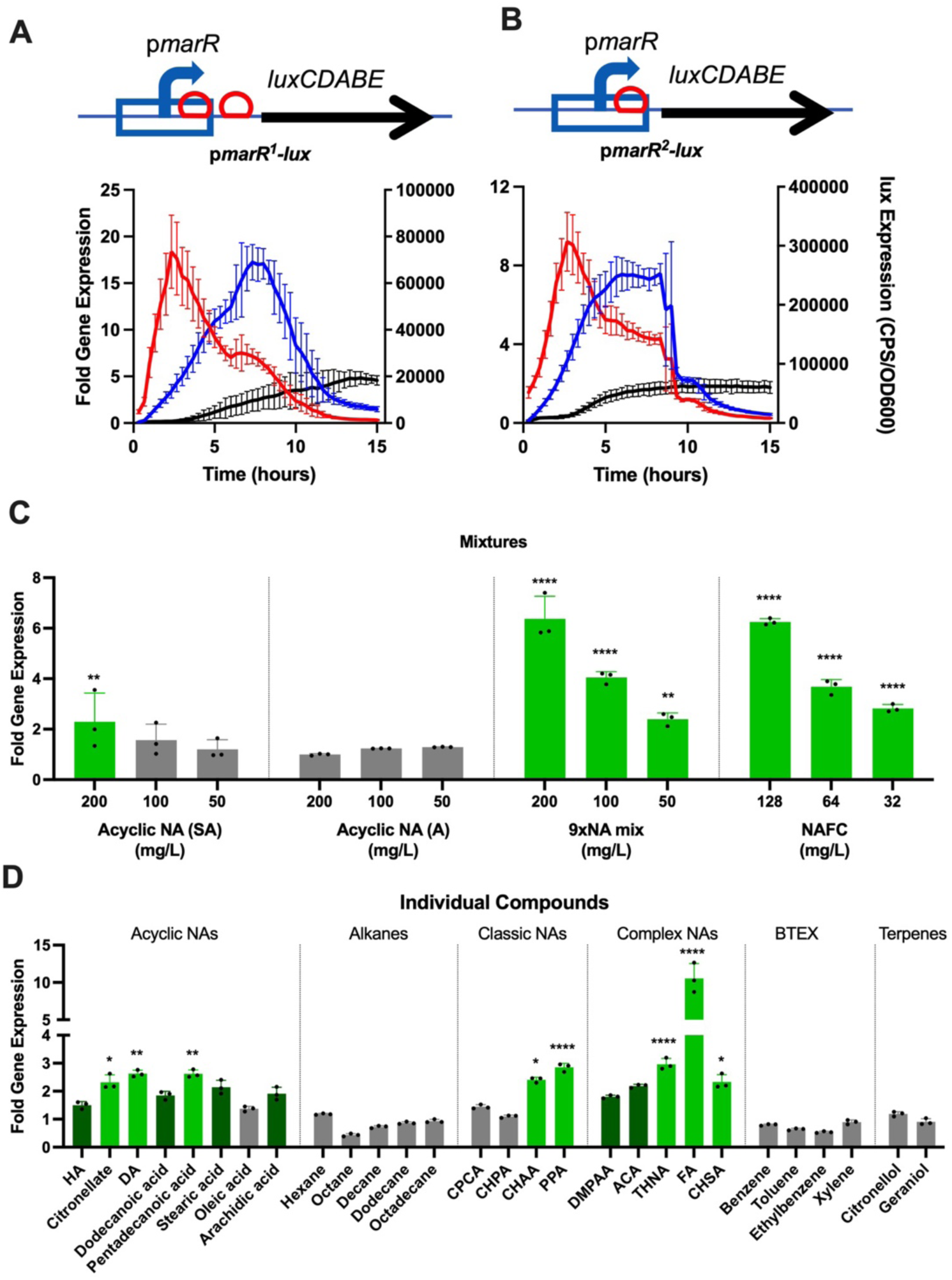
Detection of fusaric acid by first and second generation p*marR*-based biosensors and specificity of naphthenic acid detection. Gene maps of **A**) first and **B)** second generation p*marR* biosensor are shown above their respective gene expression *lux* response. Biosensors were exposed to 25 mg/L fusaric acid and *lux* expression over time was compared to untreated media controls, shown in blue and black, respectively. Fold gene expression values are shown in red. Values shown are the average and standard deviation of triplicates, and each experiment was performed at least 3 times. **C)** Fold gene expression values from the marR-L biosensor of four different naphthenic acid mixes tested across a range of concentrations. SA (Sigma-Aldrich) and A (Acros) indicate two sources of acyclic NA mixtures, the 9xNA mix and the NAFC extracts. A two-way ANOVA found that there was a statistically significant difference between the mixtures (F(4,40) = 71.49, p < 0.0001) and concentrations (F(3,40) = 91.68, p < 0.0001). **D)** Fold gene expression values of 28 different hydrocarbons at a concentration of 50 mg/L. A one-way ANOVA revealed a statistically significant difference between treatment groups (F(28,58) = 66.35, p < 0.0001). A post hoc Tukey HSD test was then conducted with each compound and mix to test significant relationships between each group (α = 0.05, n = 3). Treatment groups with significant results compared to the control are indicated with asterisks and bright green (* indicate p values below 0.05, ** indicate p values below 0.01, **** indicate p values below 0.0001). Nonsignificant groups displaying fold gene expression values above 1.5 are shown in dark green. All fold change values shown represent the average of triplicates +/− standard deviation after 3 hours of exposure and each experiment was performed 3 times.

### Specificity of naphthenic acid detection by the *marR* promoter

To identify the range of naphthenic acids compounds detected, the marR-L biosensor was screened with decreasing concentrations of four different NA mixtures, as well as 28 different individual compounds at a concentration of 50 mg/L. The *marR* promoter showed a modest induction response to high concentrations of the Sigma-Aldrich acyclic NA mixtures (**Fig 6C**), which was consistent with the RNA-seq data (p<0.05, **Table 1**). However, the *marR* promoter was induced in a dose-dependent manner to a custom mix of 9 individual NA compounds (50-200 mg/L), as well as extracted NA from oilsands process-affected water (OSPW) (32-128 mg/L) (**Fig 6C**). As fusaric acid resistance genes are induced in the presence of fusaric acid (45, 46), and in response to NAFC, this suggests the presence of antimicrobial NA compounds, or FA, within the complex NAFC mixture.

When tested for the detection of individual compounds, the *marR* promoter was specifically induced by a few acyclic NA structures, but mostly by complex and classic NA structures. The strongest inducing compound was fusaric acid (FA, 10-fold), followed by 5,6,7,8-tetrahydro-2-naphthoic acid (THNA, 3-fold), 3-phenylpropionic acid (PPA, 3-fold), cyclohexyl succinic acid (CHSA, 2.5-fold), and cyclohexane acetic acid (CHAA, 2.5-fold) (**Fig 6D**). The inducing acyclic NA included the branched compound citronellate, and longer chain compounds such as decanoic acid and pentadecanoic acid (**Fig 6D**). In summary, the *marR*-based biosensor detected diverse and complex NA mixtures and compounds and was the only sensor to strongly respond to naphthenic acids extracted from the OSPW from oil sands mining tailings ponds.

### A hypothetical gene promoter for detecting simple naphthenic acids

The promoter for a hypothetical gene at the locus tag IH404_RS03680 (referred to subsequently as ‘3680’), was upregulated in the presence of the custom NA mixture from the RNA-seq analysis (**Table S3**). While there are few identifiable functional domains in the 3680 protein, it does have three n-myristoylation sites, indicating post-translational modification with the 14-carbon unsaturated fatty acid, myristic acid. Further evidence of a potential role in fatty acid biology is suggested by the adjacent TetR regulator, a homolog of *Pseudomonas aeruginosa* DesT that detects the ratios of unsaturated to saturated fatty acids and regulates the production of unsaturated fatty acids for membrane lipid synthesis (47). We also identified a mini-Tn5-*lux* insertion mutant within this gene (*hyp3680::lux*), that also was strongly induced in the presence of the simple NA mixture (**Fig 7A**).

**Figure 7.**
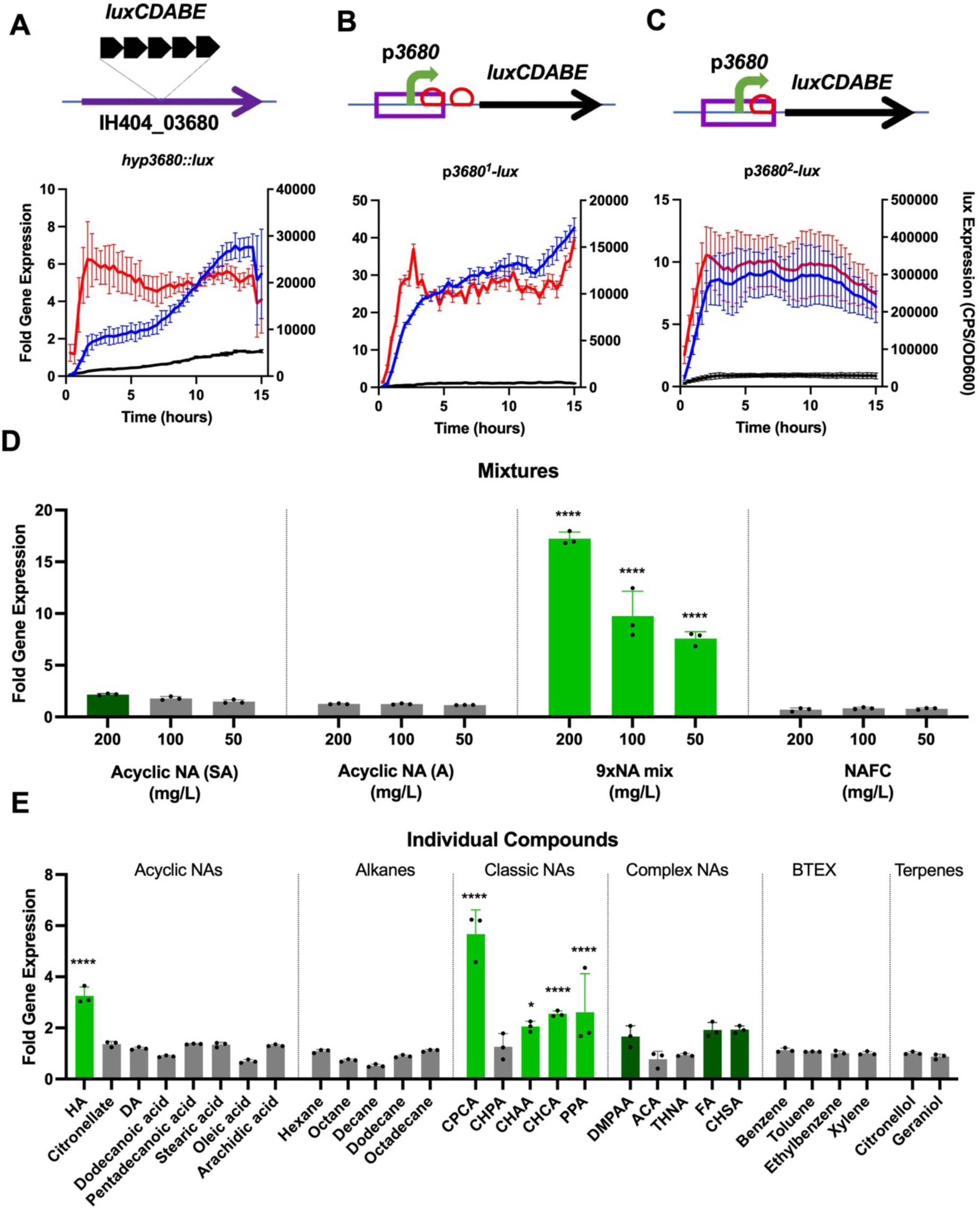
Detection of the simple 9x NA mixture with p*3680*-based biosensors and specificity to individual naphthenic acid compounds. Gene expression response from **A)** chromosomal *hyp3680::lux* reporter **B)** first generation *p3680^1^-lux* biosensor **C)** second generation *p3680^2^-lux* biosensor after exposure to 50 mg/L of the simple 9xNA mix (blue) and compared to untreated media controls (black). Fold gene expression values are shown in orange. Gene maps for each biosensor are shown above their respective *lux* response. All values shown are the average and standard deviation of triplicates and each experiment was performed at least 3 times. **D)** Fold gene expression values of four different naphthenic acid mixes with decreasing concentrations. SA (Sigma-Aldrich) and A (Acros) indicate two sources of acyclic NA mixtures. A two-way ANOVA found that there was a statistically significant difference between the mixtures (F(3,32) = 427.9, p < 0.0001) and concentrations (F(3,32) = 92.50, p < 0.0001). **E)** Fold gene expression values of 28 different hydrocarbons at a concentration of 50 mg/L. A one-way ANOVA revealed a statistically significant difference between treatment groups (F(28,58) = 21.66, p < 0.0001). A post hoc Tukey HSD test was then conducted with each compound and mix to test significant relationships between each group (α = 0.05, n = 3). Treatment groups with significant results compared to the control are indicated with asterisks and bright green (*** indicate p values below 0.001, **** indicate p values below 0.0001). Nonsignificant groups displaying fold gene expression values above 2 are shown in dark green. All fold change values shown represent the average of triplicates +/− standard deviation and each experiment was performed 3 times.

Based on the gene expression responses and a potential role in fatty acid biology, this promoter was prioritized for synthesis and construction as a plasmid based p*3680*-*lux* biosensor candidate. The chromosomal and plasmid-based p*3680*-based biosensors were all strongly induced during growth in the presence of 50 mg/L of the simple 9xNA mix (**Fig 7A-C**). The second-generation *3680* sensor exhibited almost 10-times the *lux* expression of all constructs (**Fig 7B,C**), consistent with the *atuA*, *atuR* and *marR* expression patterns (**Figs 3**, **6**). However, the first-generation *p3680^1^-lux* sensor displayed the highest fold gene expression values (**Fig 7B**) and was therefore the most sensitive to NA detection.

### Specificity of naphthenic acid detection by the hypothetical *3680* promoter

The *p3680^2^-lux* biosensor responded in a dose-dependent manner primarily to the custom mixture of 9xNA compounds, but not to OSPW extracts or acyclic NA (**Fig 7D**). After testing each separate compound from the 9xNA Mix, the *p3680^2^-lux* biosensor responded mainly to the “classic” NA, such as cyclopentane carboxylic acid (CPCA, 6-fold), cyclohexane carboxylic acid (CHCA, 3 fold), phenylpropionic acid (PPA, 3-fold) and cyclohexane hexanoic acid (CHAA, 2 fold). Additionally, a significant 3-fold *lux* response was seen with hexanoic acid (HA, 3-fold), a short acyclic NA that is also present in the custom mixture (**Fig 7E**). This sensor also responds weakly (~ 2-fold) to other more complex NA compounds such as 4-dimethylphenylacetic acid (DMPAA) and two compounds absent from the mix, cyclohexane succinic acid (CHSA) and FA.

### Dose response assays and limits of detection of the *atuA*, *atuR*, *marR* and *3680* promoters

The third-generation biosensor constructs were used to determine the sensitivity and specificity of the *atuA*, *atuR* and *marR* promoters. These complete biosensor constructs consist of the NA-inducible promoter fused to the *lux* reporter, with the inclusion of the corresponding repressor protein, under control of a “Low” strength, constitutive promoter. The atuA-L construct exhibits a linear relationship between the fold expression change values ranging from 7.8 – 125 mg/L, while the atuR-L construct displays this linear range of NA response and detection between 62.5 – 500 mg/L. The *atuR* promoter responds more slowly, with lower fold changes and less sensitivity to acyclic NA mixtures (**Fig 8A,B**). The limit of sensitivity for these promoters was determined by the NA concentration that resulted in a ~2-fold gene induction (biologically relevant) and where increasing the concentration also increased the gene expression. The limits of sensitivity were estimated to be 15 mg/L for the *atuA* promoter, and 125 mg/L for the *atuR* promoter, in response to acyclic NA.

**Figure 8.**
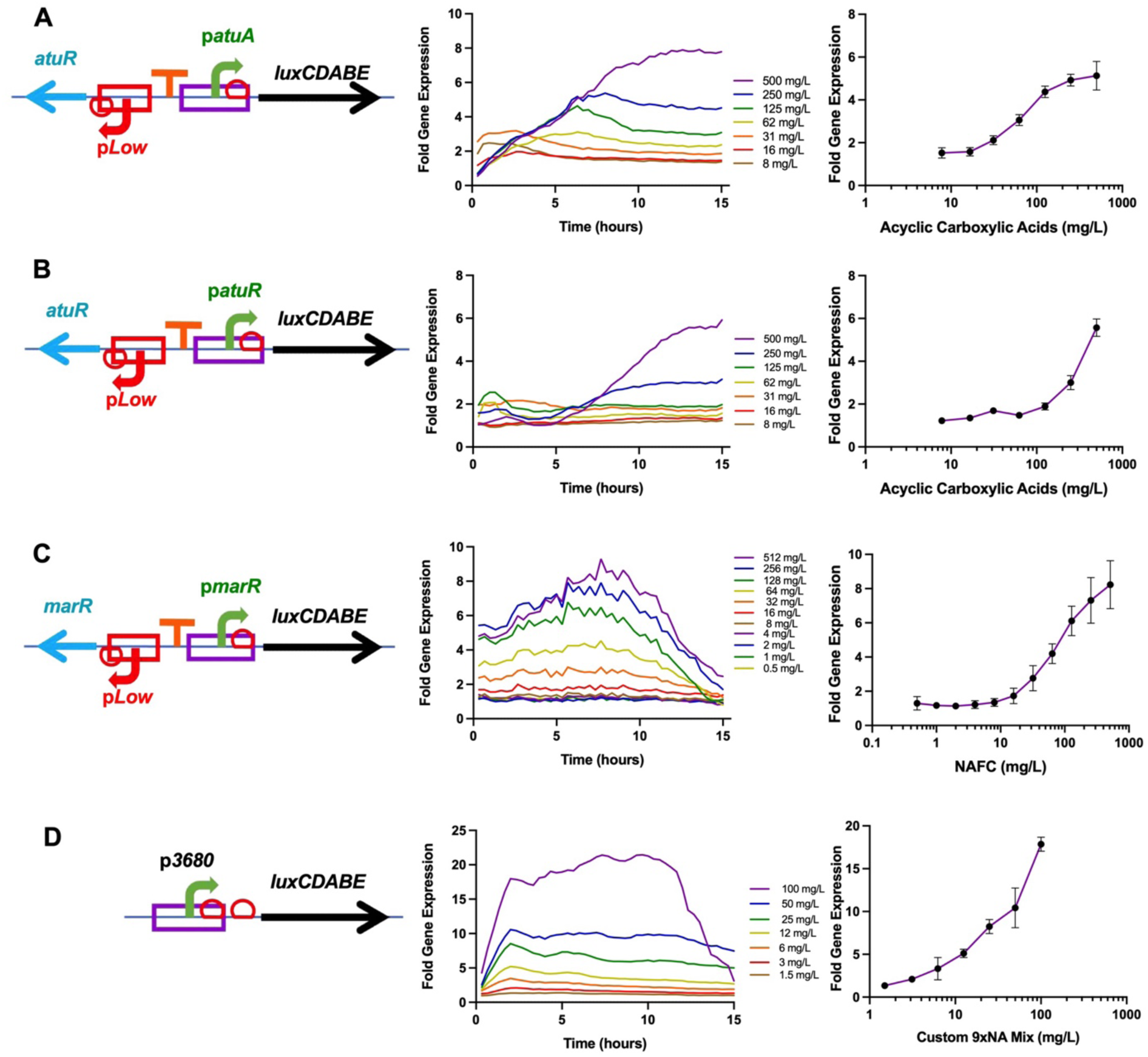
Dose responses and sensitivity of the atuA-L, atuR-L, marR-L and *p3680^2^-lux* biosensors to various NA mixtures. Gene maps of each biosensor are displayed on the left, gene expression in varying NA concentrations is shown in the middle, and dose-response curves are on the right. **A)** Fold gene expression values of atuA-L were calculated after exposure to increasing concentrations of acyclic NA, from 8 mg/L to 500 mg/L. **B)** Fold gene expression values of atuR-L were calculated after exposure to increasing concentrations of acyclic NA, from 8 mg/L to 500 mg/L. **C)** Fold gene expression values of marR-L were calculated after exposure to increasing concentrations of NAFC, from 0.5 mg/L to 512 mg/L. **D)** Fold gene expression values of *p3680^2^-lux* were calculated after exposure to increasing concentrations of the simple, 9xNA mix. All values shown are the average of triplicate experiments +/− the standard deviation, and each experiment was performed 3 times.

The third generation marR-L construct (*pLow-MarR, pmarR^2^-lux*) was screened against 12 concentrations of the OSPW mixture ranging from 0.5 to 512 mg/L and the resulting bioluminescence measurements are shown in **Figure 8C**. The *marR* promoter responded to NAFC at concentrations as low as 16 mg/L, then responded in a linear dose response as the concentration increased to 512 mg/L. The second generation *p3680^2^-lux* biosensor construct was screened with 8 concentrations of the 9xNA mixture, ranging from 1.5 - 100 mg/L (**Fig 8D**). This promoter responds very rapidly to the custom NA mixture and exhibits a strong linear, dose-dependent bioluminescent response, where the limit of detection of this custom 9XNA mix was as low 1.5 mg/L. (**Fig 8D**).

### Detection of naphthenic acids in OSPW samples using a panel of three NA-inducible promoters

In addition to demonstrating biosensor detection of individual NA compounds and concentrated mixtures, it is important to also determine the ability of these whole-cell biosensors to detect naphthenic acids in actual samples of oil sands process-affected water (OSPW). We tested the NA biosensors for their ability to detect naphthenic acids in a panel of 24 oil sands process-affected water samples, as well as NAFC that were extracted from each respective OSPW sample. Given the wide range of NAFC concentrations within the OSPW extracts (**Table S1)**, we diluted these samples by a factor of 10 or 100 in order to test a final NA concentration between 20 and 60 mg/L. In **Figure 9A**, 11/24 NAFC extracts induced a significant biosensor response above background expression levels (> 1-fold), from at least one of three whole cell biosensors. Each biosensor in this collection responds to specific NA compounds (**Figs 4-8**), which allows the estimation of abundant NA compounds present within these OSPW samples.

**Figure 9.**
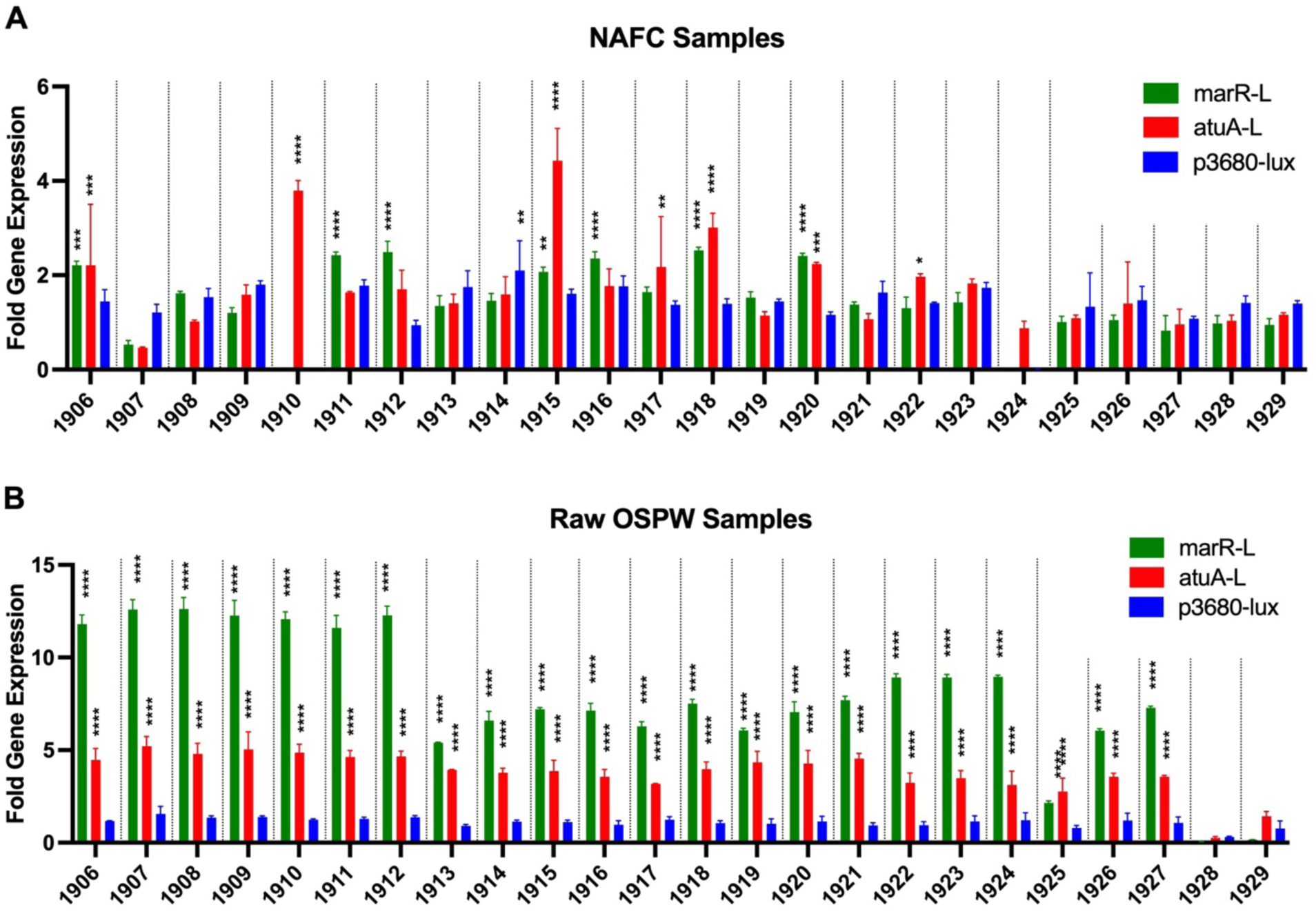
Specificity of marR-L, atuA-L, and *p3680^2^-lux* to NAFC extracts and raw OSPW samples. Fold gene expression values of marR-L, atuA-L, and the second generation *p3680^2^-lux* biosensor were taken at 8 hours, 7 hours, and 3 hours, respectively. **A.** Biosensor response to ~40 mg/L of each sample of NAFC extracts from the OSPW. A two-way ANOVA found a statistically significant difference between biosensor strains (F(2,150) = 27.85, p < 0.0001) and NAFC samples (F(24,150) = 25.41, p < 0.0001). **B.** Biosensor response to the oil sands process-affected water samples. A two-way ANOVA found a statistically significant difference between biosensor strains (F(2,150) = 5867, p < 0.0001) and raw OSPW samples (F(24,150) = 194.0, p < 0.0001). A post hoc Tukey HSD test was then conducted with each water sample and extract to test significant relationships between each group (α = 0.05, n = 3). Treatment groups with significant results compared to the control are indicated with asterisks (*p < 0.05, **p < 0.01, ***p < 0.001, ****p < 0.0001).

For example, samples 1911, 1912, and 1916 resulted in a ~2.5-fold *lux* response from marR-L, suggested that more complex multi-ringed or phenolic compounds are abundant in these mixtures. The *lux* response of atuA-L to the samples 1910 (~3.5-fold), 1917 (~2-fold), and 1922 (~2-fold), suggested that these samples have a significant proportion of acyclic NA compounds. NAFC extracted from OSPW are complex, and not surprisingly, samples 1906, 1915, 1918, and 1920 induced a bioluminescent response multiple sensors, indicating the presence complex and acyclic NA. The average 2-3-fold *lux* response of both atuA-L and marR-L to the NAFC extracts screened at ~40 mg/L in **Figure 9A** is similar to what would be expected from the standard curves (**Fig 8**).

Due to the low NA concentrations reported for the water samples (**Table S1**), 90 µl of each water sample were added to each well of the assay plate, with the addition 10 µl of 10x BM2 growth media, to allow for minimal dilution of the OSPW sample. In **Figure 9B**, the biosensor panel performed much better, where 22/24 water samples induced a significant biosensor response, from at least two of three whole cell biosensors. The marR-L and atuA-L biosensors displayed a high fold expression response (between 4 and 12-fold) for all samples except two, while the promoter for *3680* did not respond to any water sample. The discrepancy between the biosensor outputs when exposed to a water samples and extracts of the same water sample could possibly be explained by a bias or inefficiency in NA extraction of tailings water samples.

## Conclusions

Whole cell bacterial biosensors have been previously constructed that are specific to various hydrocarbons, including alkanes (48, 49), benzene/BTEX (50, 51) and naphthalene (52). Here we describe a bacterial biosensor approach to construct naphthenic acid biosensors. We identified several strong inducible promoters capable of NA sensing, driving the expression of genes involved in degradation or protection from NA. A transcriptome approach was used for the discovery of NA-induced genes, although there were relatively few significantly induced genes when exposed to various NA extracts, compared to other genomic studies (53). This may have been a limitation of using duplicates for the transcriptome studies, but the use of transcriptional *lux* fusions validated many of the upregulated genes identified by RNA-seq. Many fatty acid degradation genes that likely contribute to β-oxidation of naphthenic acids were identified, which is consistent with the results of a proteomic study with *Pseudomonas fluorescens* Pf-5 (53). This study identified transcriptional repressors that may act as naphthenic acid sensing proteins. The TetR and MarR family of transcription factors are well characterized to sense and respond to small molecules, and in turn regulating a wide variety of cellular processes including multidrug efflux pumps and carbon utilization. For these reasons, small molecule sensing repressor proteins are commonly used in the construction of bacterial biosensors (11, 48, 54).

The NA biosensors in this study can sense and respond to different NA mixtures with good sensitivity, and each promoter demonstrates a unique naphthenic acid specificity profile. The NA-induced promoters are strongly induced in a concentration-dependent manner, which may allow for estimation of NA concentrations in OSPW by extrapolation from the standard curve of gene expression responses (**Fig 8**). When used to detect NA in small volume samples of water taken from tailings ponds, there was NA detection by 2/3 biosensors in almost all water samples. When NAFC of those corresponding water samples were tested at ~40 mg/L, fewer samples were detected, which may be due to a change in NA composition after organic extraction. The ability to test small volumes of raw water samples provides another advantage of minimal sample preparation. Naphthenic acid biosensors show great potential for a rapid, reliable, cost-effective, and semi-quantitative method for detection of environmental pollutants.

## Acknowledgements

The authors acknowledge Laetitia Mazenod and Rich Moore for technical assistance, and Paul Gordon for bioinformatics support. Funding was provided by an NSERC Discovery Grant and Mitacs Canada.

## Notes

### Competing Interest Statement

The authors have declared no competing interest.

